# Linking macroscale structure and function in brain-like recurrent neural networks

**DOI:** 10.64898/2026.02.19.706880

**Authors:** Peiyu Chen, Zaixu Cui, Christos Constantinidis

**Affiliations:** Department of Biomedical Engineering, Vanderbilt University, Nashville, TN 37235; Chinese Institute for Brain Research, Beijing 102206, China; Program in Neuroscience, Vanderbilt University, Nashville, TN 37235; Department of Ophthalmology and Visual Sciences, Vanderbilt University Medical Center, Nashville, TN 37212

**Author notes:** Correspondence (P.C.).

## Abstract

Linking structure and function is a central topic in both neurobiology and artificial intelligence. Human brain functions are organized across the macroscale cortex into parcellations, modules, and hierarchies that can be inferred from intrinsic structural architecture, providing an interpretable and clinically meaningful framework for linking structure to function. While artificial neural networks have been successfully aligned with human cognition at the representational level, it remains unclear whether the structural principles linking brain anatomy and function can extend to artificial neural networks, and whether imposing brain-like structural constraints can induce comparable functional organization to that observed in the human brain. Here, we introduce BrainRNN, a brain-like recurrent neural network architecture inspired by macroscale human cortical structure. We show that under structural constraints, BrainRNNs selectively regulate the distribution of connectivity and recruit more activated units in association regions for higher-order cognitive capacity. Moreover, we demonstrate structure–function coupling in BrainRNNs and show that structural constraints enable macroscale functional organization, including functional modules and gradients, to emerge along topographic and topological axes, closely mirroring empirical findings in the human cortex. Together, these results demonstrate how multiple brain-like structural constraints jointly shape functional organization and enable function to be inferred from structure, highlighting the potential of structurally grounded artificial intelligence for neuroscientific research.

## Introduction

The relationship between structure and function is a central problem in both neurobiology and artificial intelligence. In the human brain, macroscale functional organization, including functional parcellations, modules, and hierarchies, has been widely characterized in relation to anatomical structure^1–6^. These organizational principles provide an intuitive and clinically meaningful framework for linking brain anatomy to cognitive function and behavior^7–9^. Importantly, such functional organization can be inferred from structural architecture and intrinsic connectivity patterns in the absence of explicit task activation^3,6^. Despite striking progress in modern artificial neural networks, most interpretability approaches rely on post hoc analyses of task-evoked representations like gradient-based attribution to assign functional relevance to units^10,11^. While informative, these methods do not infer function from structure, as is commonly done in neuroscience^1,12^.

Artificial neural networks have been widely used to relate their learned computations to human cognitive function^13^. Most prior work has emphasized the alignment of functional representations with empirical neural data^14–17^, whereas their differences in underlying architecture have received far less attention^18,19^. High-performing artificial neural networks often rely on freely optimized connectivity with goal-driven training^20,21^. Few studies have incorporated structural considerations, and these have largely focused on isolated factors^18,22^ or specific functional areas^14,16,19^, leaving the perspective from the macroscale structural differences unexplored. Consequently, although artificial neural networks can produce functional representations that align with neural activity, it remains unclear whether their internal organization supports direct comparisons with canonical findings in large-scale brain functional organization.

Biological cognition emerges from neural systems that are simultaneously spatially embedded, structurally constrained, and functionally adaptive^23–26^. Different from artificial neural networks, the brain is not a generic network of interchangeable units: neurons and cortical areas are embedded in physical space^27^, organized along robust topographic manifolds^26^, and connected by anatomical pathways that are shaped by wiring economy^28^. These structural principles fundamentally limit the information flows^29,30^, organize neural representations^31,32^, and ultimately, support cognitive computations^7,12^. Recent neuroscientific research highlights both topographic and topological factors as two key structural aspects that anchor cortical function^26,33^. Topographic structure embeds neural populations in physical space, constraining the spatial distribution of neural circuits^24,34^. Topological structure, including long-range connectivity and network modularity, enables information integration across distributed regions^1,28,35^. These structural bases jointly shape the pattern of functional activations and further give rise to large-scale functional organization across the cortex^2,26,36^. Despite extensive correlational evidence, the causal role of macroscale structural constraints in shaping functional organization remains poorly understood. Moreover, it is unclear whether similar organizational principles can emerge in artificial recurrent networks, either with or without explicitly imposed structural constraints.

To fill these gaps, we proposed a brain-like recurrent neural network (BrainRNN) that embeds units within a hemispheric coordinate system, incorporates area-specific visual input and motor output interfaces inspired by cortical organization, and is trained under wiring-cost constraints. This design imposes biologically motivated topographic and topological constraints on the circuits that support a broad range of cognitive tasks. We hypothesized that structural constraints shape function and, in turn, allow functional organization to be inferred from the structure, as is commonly done in systems neuroscience. Specifically, we first examined how structural constraints in the BrainRNNs shape connectivity, functional activation patterns, and task performance. Second, we evaluated whether the trained BrainRNNs exhibit structure–function coupling comparable to the cerebral cortex. Third, we evaluated whether functional organization, specifically functional modules and gradients, emerges from both the topographic and topological structure in the BrainRNNs, as observed in the human brain, and how the organization is shaped by structural constraints. Therefore, by integrating macroscale brain-like structural constraints into the artificial neural networks, we aimed to demonstrate whether principles linking structure and function in the human brain can also be applied to artificial neural networks, and how these structural constraints can shape functional organization.

## Results

### Brain-like recurrent neural networks

We introduce “brain-like” recurrent neural networks (BrainRNNs) by embedding units into a hemisphere manifold and assigning areal interfaces that simulate primary visual and motor topographies (Fig. 1A). Recurrent units were distributed quasi-uniformly across a hemispheric manifold using a Fibonacci lattice^37^, with each unit anchored to a fixed location in three-dimensional space along the left–right (L–R), posterior–anterior (P–A), and dorsal–ventral (D–V) axes of a hemisphere, analogous to the cortex. Areal interfaces with external inputs and outputs are critical for macroscale brain organization, but have long been underestimated. External visual input was imposed on visual units in the posterior region, aligned with the primary visual cortex. Motor readouts were taken from motor units in an anterior zone positioned slightly rostral to the midline, analogous to the primary motor cortex. Other units, located outside the visual and motor regions, were defined as association units. This geometric arrangement enforces a flow of information along the topographic regions: visual signals enter through posterior units, undergo recurrent processing, and are subsequently decoded by anterior motor readout units to achieve the task goal.

**Fig. 1.**
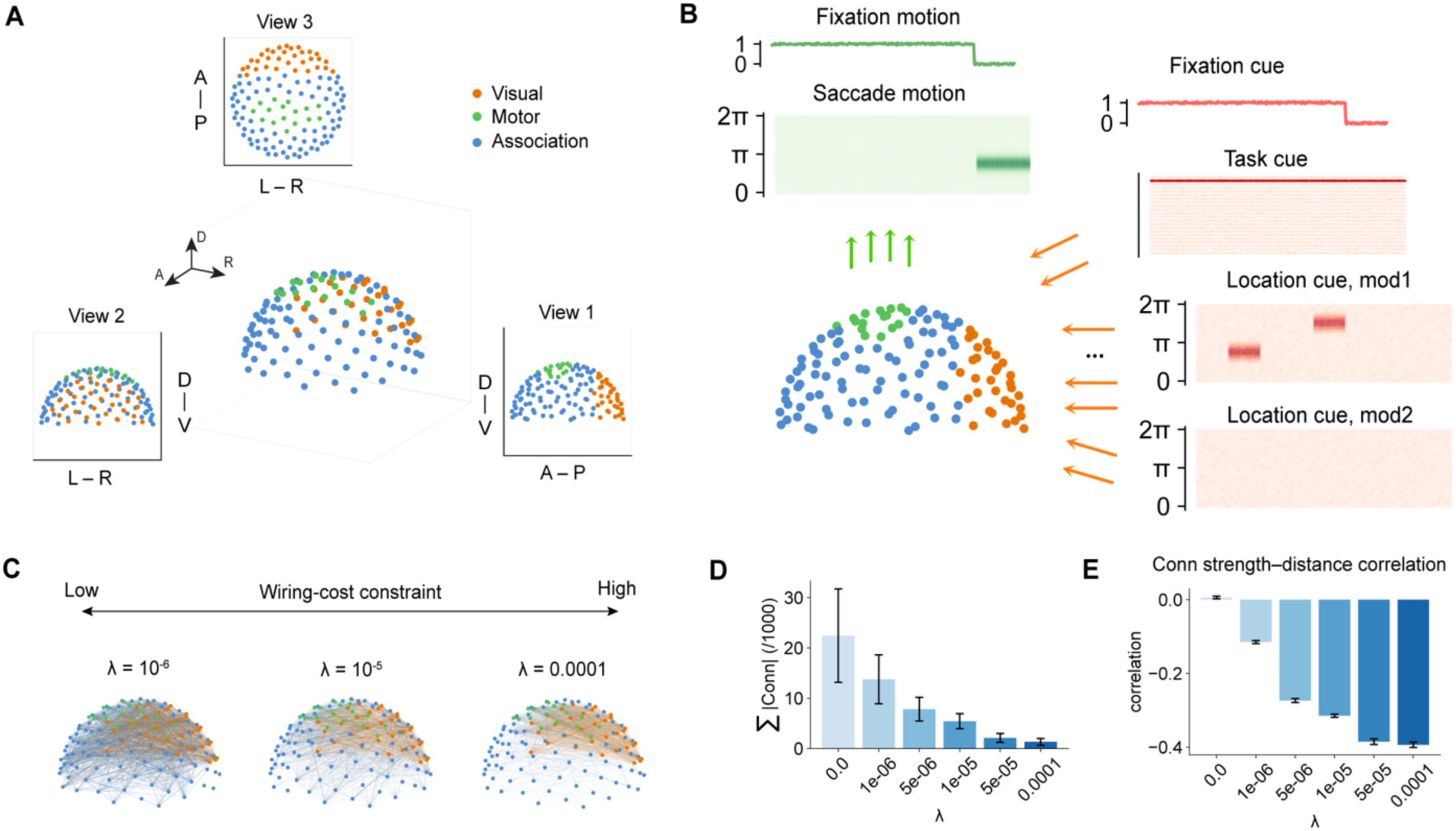
| BrainRNN architecture and training paradigm. **A**, A brain-like recurrent neural network (BrainRNN) with 128 recurrent units embedded on a cortical-like surface partitioned into visual (32 units, orange), motor (16 units, green), and association (80 units, blue) territories. Renderings are shown from right (bottom right, View1), anterior (bottom left, View2), and dorsal (top, View3) viewpoints. **B**, Input (visual) and output (motor) schematic for the Remember-first task (viewed from the right side). Visual inputs (fixation cue, task cue, and stimulus locations of two modalities; orange) are passed through an input layer to target visual units. Motor outputs (fixation and saccadic location; green) are read out from motor units through an output layer. **C**, 3-Dimensional view of the BrainRNNs trained under different wiring-cost constraint parameters λ. The width of the line represents the connection strength (absolute value of connection weights). Top 10% absolute value of connection weights are displayed here. Connections within visual, motor, and association territories are colored orange, green, and blue, respectively; all other connections are colored in gray. **D**, Mean total connection strength across BrainRNNs trained with different wiring-cost constraints parameters λ. **E**, Correlation between inter-unit distance and the connection strength for each wiring-cost constraints parameters λ. Abbreviate: A-P: anterior-posterior; L-R: left-right; D-V: dorsal-ventral; |Conn|: connection strength.

BrainRNNs were trained to perform twenty-two cognitive tasks spanning four task families: visuomotor (VM), working memory (WM), match–nonmatch (MNM), and decision-making (DM) (see Methods, Extended Data Fig. 1, and Supplementary Table 1 for detailed task definitions). For each task, the network received three types of input through the visual units: a fixation cue, a task cue, and spatial location cues (Fig. 1B for the coding of model and Fig. S1A for an example task scene). Fixation and saccade directions were read out from the motor units. For example, in the Remember-First working memory task, BrainRNNs received two spatial locations during the stimulus period during two stimulus periods but were required to remember the first location and saccade to that location during the response period (Fig. 1B). We trained BrainRNNs with 128 recurrent units each to perform all 22 tasks, for a maximum of 300,000 training sessions. Training was stopped early once all tasks achieved an accuracy greater than 0.95 (see Fig. S1B for an example accuracy curve during training).

Interareal distance is a key determinant of anatomical structural networks in the brain, which is demonstrated to obey an economical connectivity pattern^28^. Structural connectivity exhibits a distance-dependent decay, such that the probability of connections decreases with increasing interareal distance, reflecting a trade-off between wiring cost and functional integration. To capture this biological trade-off^28,35^, we incorporated a wiring-cost regularization term with the parameter λ into the training objective^18^. Previous studies highlight that this regularization equips trained neural networks with topological characteristic akin to a brain network^18^. Specifically, we penalized recurrent weights multiplied by the corresponding inter-unit distance, determined by the pre-defined unit coordinate on the hemisphere, using a L1 regularization (Fig. 1B). This constraint discourages strong and long connectivity, thereby reducing wiring resource usage^35^. As expected, the sum of connection strengths (absolute values of connectivity weights) decreases with increasing wiring-cost constraint parameters, providing a more economical wiring solution (Fig. 1C). In addition, we calculated the correlation between the inter-unit distance and connection strengths (Fig. 1D). Wiring-cost regularization induced a negative correlation between distance and connectivity strength (absolute value of connection weights), and increasing the wiring-cost penalty led to stronger negative correlations, consistent with economical wiring principles observed in brain networks^35^. Taken together, the topographic embedding and wiring-cost economy prior impose anatomical constraints on the network. These constraints ensure that information transformations from sensory input units to motor output units emerge through spatially structured recurrent dynamics, providing a referential network that mirrors key aspects of macroscale human brain anatomy and supports sensorimotor cognitive tasks.

### Structural constraints sculpt functional activations and cognitive capacity

Structural constraints anchor functional activation patterns and thereby shape the behavior. In conventional RNNs without structural constraints, external inputs can reach all neurons directly, and recurrent connections are effectively cost-free, different from the neurobiological condition. By contrast, BrainRNNs incorporate multiple structural factors observed in the biological brain, which could anchor the flow of information. To assess how such structural constraints shape network function, we first quantified the distribution of activated units across all tasks. Because BrainRNNs use rectified linear unit (ReLU) activations, some units can remain inactivated throughout processing. A unit in the BrainRNN was therefore defined as activated if it exhibited nonzero activity.

We found that BrainRNNs trained with different wiring-cost constraint parameters λ developed distinct selection of connectivity types, which gave rise to distinct activation maps. As the wiring-cost parameter λ increased, BrainRNNs exhibited a monotonic decrease in the total number of activated units (Fig. 2A–B). Notably, the number of activated visual and motor units remained relatively stable across values of λ, whereas the number of activated association units decreased sharply. Under stricter spatial constraints (λ = 5 × 10⁻⁵ or 10⁻⁴), nearly all activated units were confined to the visual and motor regions, with only a small number of association units located in the intermediate regions between them. Few or no activated units were observed in other distant areas from the visual and motor units, particularly in anterior regions. Thus, the average distance of activated units decreased with increasing λ (Fig. S2A). To examine how network structure supports these distinct activation patterns, we quantified the proportion of recurrent connection strength associated with visual, motor, and association units, defined as the sum of connection strengths linked to units in each group divided by the total recurrent connection strength (Fig. 2C). As wiring-cost constraints increased, the proportion of association connection strengths decreased from 82.3% to 16.7% while the proportion of visual connection strengths increased from 37.7% to 76.3% and the proportion of motor connection strengths increased from 20.6% to 27.5%. This connection change can be observed in Fig. 1C. This shift in connectivity composition explains the differential spatial distribution of activated units across models trained with varying wiring-cost constraints.

**Fig. 2.**
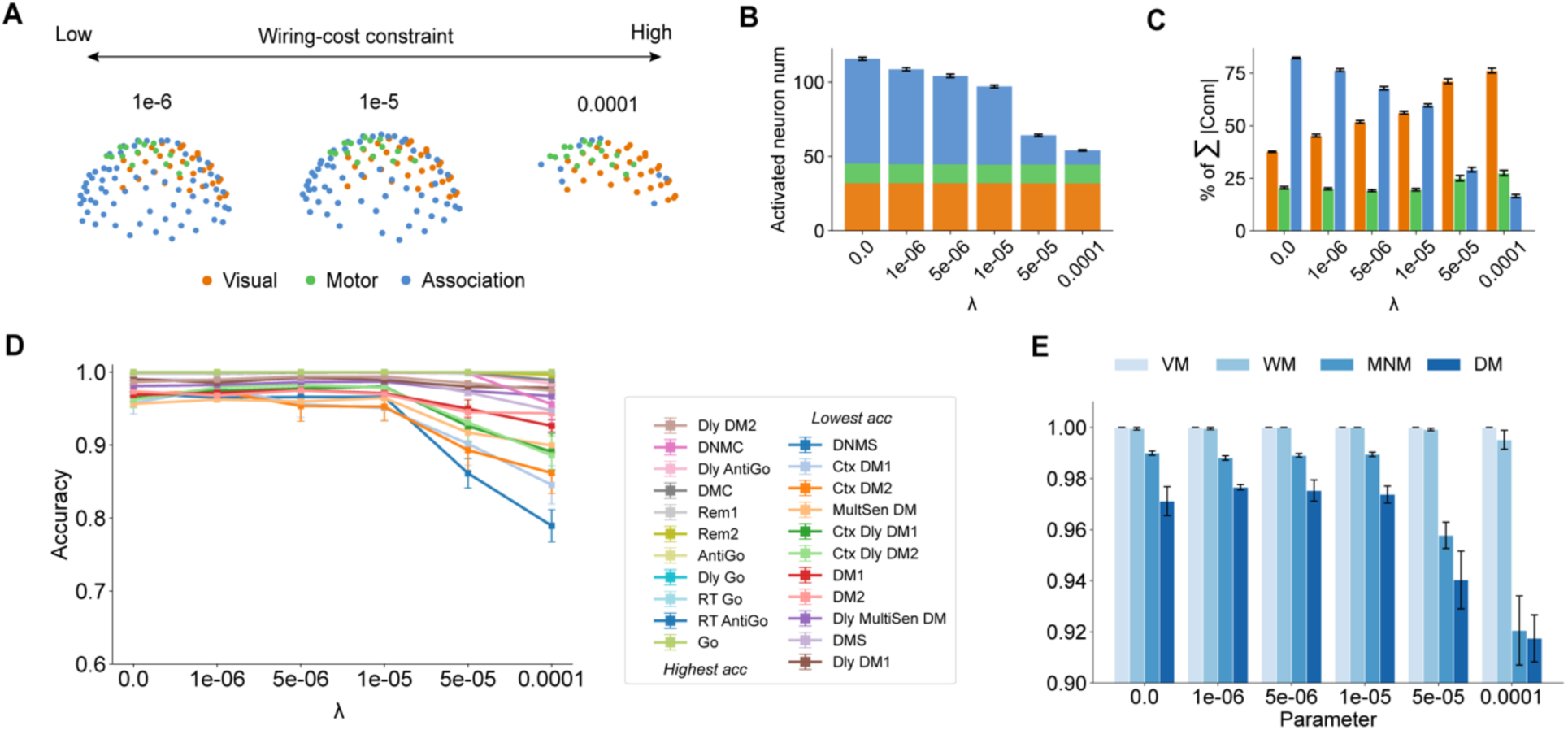
| Structure constraints regulate activation and cognitive capacity. **A**, Distinct post-training activation map across different wiring-cost constraint parameters. As Fig. 1A, units are attributed to the visual (orange), motor (green), or association (blue) regions. **B,** Increasing the wiring-cost penalty (λ) decreases the number of activated units, mainly in the association regions. **C,** Proportion of recurrent connection strengths associated with visual, motor, and association units. **D,** Task accuracy after training across λ. Task names in the legend are ranked by average accuracy across all models. **E**, Family-level average task accuracies across models trained with different λ. Four task families are visuomotor (VM), working memory (WM), match–nonmatch (MNM), and decision-making (DM). Individual tasks are defined in Methods. Error bars in **D–E** denote SEM.

We hypothesized areal interfaces of visual input and motor output, important characteristics of large-scale brain organization, play a key role in shaping the activation pattern. To test this, we conducted an ablation experiment. Firstly, the areal interfaces for visual input and motor output were removed, such that any unit could receive external input and be freely read out. All other settings were kept identical, including the same hemispheric coordinate presets and wiring-cost constraints, so we named it as Conventional RNNs (CRNNs; when λ= 0, no structural constraints) or Spatial-embedded RNNs^18^ (SpRNNs; when λ > 0, only hemispheric spatial embedded). As a result, although the overall sum of connection strengths (three-dimensional views in Fig.S2B) decreases with increasing wiring-cost parameters λ (Fig. S2C), all units were activated, even when wiring-cost constraints were applied during training (Fig. S2D). In the absence of areal interfaces of visual input and motor output, any unit could receive external input and be freely read out, resulting in unconstrained information flow to activate all units. Secondly, in the BrainRNN, the visual and motor units are less distributed than association units, so the activation pattern differences may be just resulted from their inter-unit distances with wiring-cost constraints. To reject this assumption, we randomized the spatial locations of visual, motor, and association units in the BrainRNNs and repeated the same analyses (Fig. S2E–J). Under this manipulation, the input and output interface was no longer spatially localized but randomly distributed across the hemisphere. Despite this disruption, selective activation patterns still emerged as λ varied, with less change than the BrainRNN in Fig. 2C. For example, as λ increased, the proportion of association connection strengths decreased from 83.3% to 38.2%. Unlike the original BrainRNNs (Fig. 2A), the activated units were no longer spatially clustered but instead distributed randomly across the hemisphere as λ increased (Fig. S2F). Together, these results indicate that the spatially organized activation patterns observed in BrainRNNs do not arise solely from predefined spatial coordinates, but critically depend on the structured input (visual) and output (motor) interfaces, which constrain and shape functional activation during training.

We next evaluated whether these activation and connection differences under different wiring-cost parameters λ were reflected in task performance. We assessed performance on individual tasks (Fig. 2D; task names in the figure legend are ranked by average accuracy across all models) and the average performance within each task family (Fig. 2E). BrainRNNs maintained near-ceiling performance on the simplest visuomotor tasks, even when the population of activated units was small and largely confined to visual and motor regions under the strongest wiring-cost constraint (λ = 10⁻⁴). By contrast, performance on tasks in the match–nonmatch and decision-making families declined markedly once λ ≥ 5 × 10⁻⁵, coinciding with the sharp reduction in the number of activated association units. The largest performance decrements were observed for the delay non-match-to-sample (DNMS) and contextual decision-making (Ctx DM) tasks, which place heavier demands on information maintenance and integration. Together, these results suggest a cognitive hierarchy of task performance, ranging from simple visuomotor task to the more demanding match–nonmatch and decision-making tasks, which require broader recruitment of activated units in the association region. Our results imply that under specific structural constraints, activation and connections in the association region can be linked to cognitive capacity. This pattern parallels functional organization in the brain, where association cortices play a central role in higher-order cognition^4^.

In summary, BrainRNNs provide a brain-like framework for understanding how structural constraints influence function. We found that structural constraints regulate the distribution of connections across different regions (visual, motor, and association), which anchors both the number and spatial distribution of activated units. Under structural constraints, the expansion of activated association units is associated with increased higher-order cognitive capacity during demanding tasks. More broadly, these findings offer a mechanistic link between the expansion of association regions located far from primary sensory and motor cortices (such as the prefrontal cortex in the anterior regions^4^), structural constraints on neural wiring, and the functional demands of higher-order cognition.

In the following section, we focus on two classes of structural constraints widely studied in the macroscale brain organization. We define here (i) topographic structure, as the pre-specified hemispheric coordinates of units with different functions (visual, motor, and association); and (ii) topological structure, as the recurrent connections between units, which determine their functional communication. We aimed to investigate whether the functional organization in the BrainRNNs are distributed along both topographical and topological axes by evaluating the structure-function coupling, functional modules, and functional gradients, three key discoveries in the human brain.

Moreover, the wiring-cost constraint provides a key mechanism through which topographic organization is embedded into network topology (Fig. 1D&E) and subsequently shapes activation patterns and functional properties (Fig. 2). By varying the strength of the wiring-cost parameter, we systematically vary the degree to which structural constraints are imposed on the BrainRNNs. Prior studies demonstrated that such constraints promote the locally compact functional activations^18,19^, and can improve the prediction empirical neural activity^19^. Here, we conduct a more comprehensive analysis to investigate how wiring-cost constraints shape functional organization and enhance the interpretability of function from the structure.

### Structure–function coupling in the BrainRNNs

The brain functional activity is shown to be anchored by both topographic structure (like spatial distance) and topologic structure (like white-matter anatomical tracts), a relationship commonly referred to as structure–function coupling^27,29,36^. We evaluated whether BrainRNNs exhibit similar structure–function relationships. As a reference, we used multimodal human brain MRI data from the Human Connectome Project (HCP)^38^. Details regarding participant demographics, selection criteria, and preprocessing pipelines are provided in the Methods. The group-averaged Euclidean distance matrix, structural connectivity matrix (white-matter tracts), and functional connectivity matrix (Pearson correlation coefficient of temporal resting state fMRI signals), were constructed based on the Schaefer-400 template with 400 atlases and each be described as a 400×400 matrix (Fig. 3A)^3^. We constructed these three matrices for BrainRNNs (Fig 3B). The distance matrix was defined as the Euclidean distance between the preset coordinates of units. Structural connectivity was quantified as the recurrent connection strengths between each pair of units. Functional connectivity was computed as the Pearson correlation between the trial-averaged temporal activity of two units in the absence of task cues. Only activated units were involved in this analysis, as non-activated units do not contribute to network function.

**Fig. 3.**
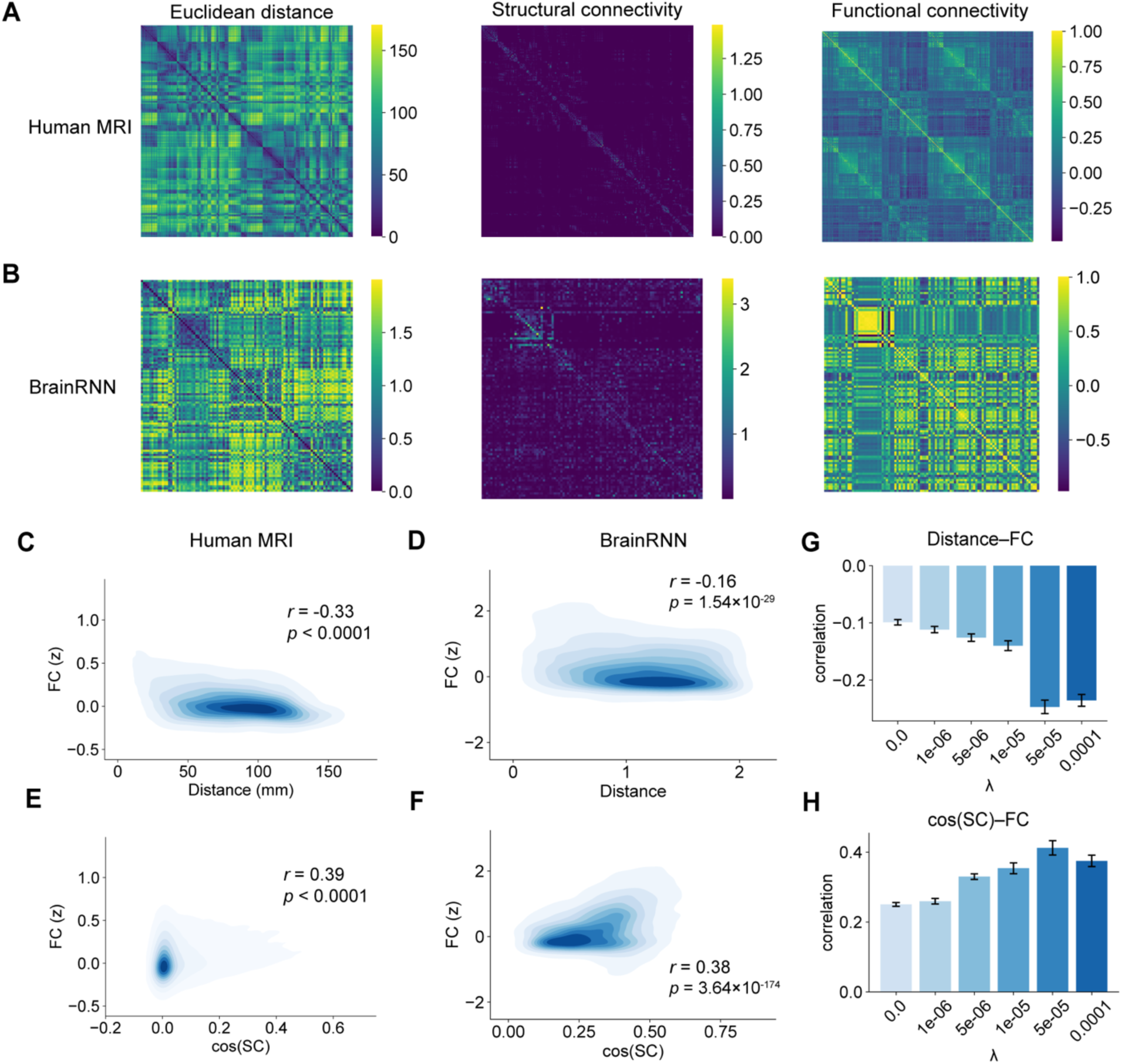
| Structure–function coupling in the brain and BrainRNNs. **A–B**, Spatial distance, structural connectivity, and functional connectivity of human MRI (**A**) and a BrainRNN model (**B**). **C–D**, Inter-unit Euclidean spatial distance is negatively correlated with functional connectivity in both Human MRI data (**C**) and the BrainRNN model (**D**). **E–F**, Cosine similarity of structural connectivity (abbreviated as cos(SC)) is positively correlated with functional connectivity in both human MRI data (**E**) and the BrainRNN model (**F**). **G–H**, Pearson correlation between Inter-unit spatial distance (**G**) or cosine similarity of structural connectivity (**H**) with functional connectivity across wiring-cost constraint parameters. Abbreviate: SC: structural connectivity, FC: functional connectivity. Error bars in **G–H** denote SEM.

The structure–function coupling is quantified as the Pearson correlation between the structural profiles (including spatial distance and structural connectivity matrices) and functional connectivity matrices^36^. Fisher’s z-transformation is applied at functional connectivity for correlation analysis. In the human brain, functional connectivity is negatively correlated with spatial distance, indicating that regions farther apart tend to exhibit more differentiated functional activity (Fig. 3C). We observed a similar negative correlation in BrainRNNs (Fig. 3D). Structural connectivity in the brain is sparse, so not all pairs of functionally connected regions are directly linked by anatomical connections. Moreover, functional correlations can reflect indirect interactions or shared connectivity with other regions. Therefore, as previous studies, we computed the cosine similarity between structural connectivity and correlates it with functional connectivity^39^. Both human MRI data and BrainRNNs exhibited a significant positive correlation between the cosine similarity of structural connectivity and functional connectivity (Fig. 3D–E).

To further examine how structural constraints shape structure–function coupling, we quantified coupling strength across all wiring-cost parameters (Fig. 3G–H). A significant negative correlation between spatial distance and functional connectivity was present in all cases (*p* < 0.0001 for all models), and its strengths increased as a function of increasing wiring-cost constraints up to λ = 5 × 10^-5^. We note that this distance-dependent functional difference was present even in the absence of a wiring-cost constraint (mean *r* = -0.11 when λ = 0). This suggests that the imposed topographic organization that included the areal visual (input) and motor (output) interfaces, can give rise to a functional differentiation between distant units without the influence of topological structure. Similarly, the cosine similarity of structure connectivity remained significantly positively correlated with functional connectivity across wiring-cost parameters (*p* < 0.0001 for all models).

By comparison, both CRNNs and SpRNNs also exhibited a positive correlation between the cosine similarity of structural connectivity and functional connectivity, but with lower correlation values than BrainRNNs trained under the same wiring-cost constraints (e.g., mean r = 0.20 for SpRNNs and mean r = 0.34 for BrainRNNs when λ = 0.0001, Fig S3A). However, they exhibited very low correlation between spatial distance and functional connectivity (e.g., mean *r* = -0.02 when λ = 0.0001, Fig. S3B). This might be because external input of SpRNNs influences more units than BrainRNNs which cannot be explained by both the structural connectivity and distance. These results indicate that, compared with CRNNs and SpRNNs, with brain-like structural constraints, BrainRNNs exhibit structure–function coupling patterns that more closely resemble those observed in the brain.

### Structure anchors functional organization, enabling the discovery of functional modules

Local differentiation and network integration are regarded as complementary principles of large-scale brain organization^1,40^. These principles give rise to functional modules that are both topographically localized and distributed across cortical regions based on wiring topology. Previous studies have primarily characterized functional modules using resting-state fMRI functional connectivity without task-evoked responses, which reflects topological architecture of intrinsic communication, as exemplified by canonical functional networks identified by Yeo et al. (Fig. 4A)^41^. These studies implied that functional modules distributed along topographic and topological structures.

**Fig. 4.**
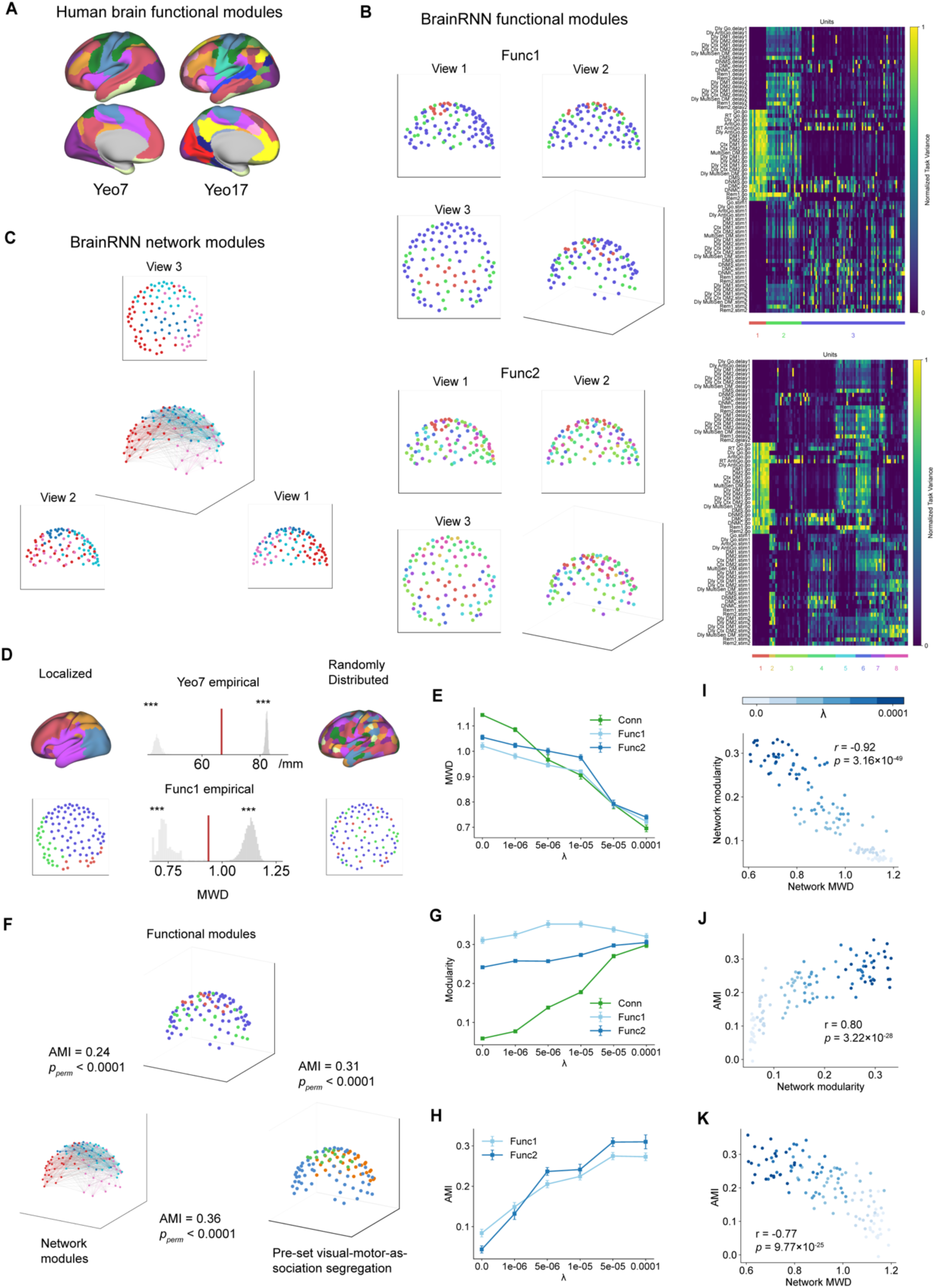
| Structure shapes functional modular organization. **A**, Macroscale human functional modules derived from the resting-state functional connectivity. Yeo7 and Yeo17 refer to the 7- and 17-modular network parcellations identified by Yeo et al.^2^. **B,** Functional modules (func1 and func2) of a BrainRNN model calculated by clustering the normalized task variance (right, units reordered by module membership). Units are displayed in their three-dimensional spatial coordinate and colored by functional module labels (left). **C,** Network modules of the BrainRNN calculated by clustering the recurrent connections. Units are colored by network module labels. Within-module connectivity is colored by their modular color while between-module connectivity is colored in gray. **D,** Functional modules of both the human brain (Yeo7) and BrainRNN (Func1) are neither purely localized nor randomly distributed (*p* < 0.001). Examples of spatially localized (left) and randomly distributed (right) functional modules are shown. The empirical mean within-module distance (MWD) of functional modules (red line) lies between the distributions of localized (light gray) and random (dark gray) modules. **E**, MWDs of both network and functional modules decrease with increasing wiring-cost constraint λ. **F**, Functional modules are anchored by network modules and predefined visual–motor–association territories, quantified using adjusted mutual information (AMI). **G**, Modularity of functional and network modules across BrainRNNs trained with different wiring-cost constraint parameters λ. **H**, AMI between functional and network modules increases with the increase of wiring-cost constraints λ. **I**-**K**, Pairwise correlations among MWDs of network modules, modularity of network modules, and AMI between functional (Func1) and network modules across BrainRNNs trained with different wiring-cost constraint parameters λ. Error bars in **E, G, H** denote SEM. *** denote *p* < 0.001.

We investigated how functional modules in BrainRNNs can be linked to topographic and topological structure by examining the relationship between functional modules and topographic coordinates, predefined visual-motor-association segregation, or network modules. In human experiments, functional responses are typically probed using a limited set of tasks or are confined to relatively restricted brain regions, which constrains the identification of complete functional modular organization. By contrast, BrainRNNs allow access to full activation patterns across all tasks, enabling the characterization of precise functional modules. The functional modules of BrainRNNs are calculated by clustering the normalized variance of all cognitive tasks across different periods (Fig. 4C for a sample model with λ = 10^-5^)^21,42^. Similar to multiscale functional modules observed in the macroscale human brain (Yeo7 for 7 clusters and Yeo17 for 17 clusters, Fig 4A)^2,3^, we calculated the optimal clustering solutions at both coarse-grained (2–5 clusters; denoted as Func1) and finer-grained clustering (6–30 clusters; denoted as Func2) of the normalized task variance (Fig 4B). Network modules of the structural connectivity were calculated using the Louvain algorithm (Fig. 4C for BrainRNNs and Fig. S4A using structural connectivity in the Human MRI)^43^.

We evaluated whether BrainRNNs exhibit functional modules that are both spatially localized and distributed along topographic coordinates, as observed in the brain^1,40^. We quantified the level of spatial compactness using the mean within-module distance (MWD). Two null distributions representing absolute localized and randomly distributed were constructed to assess statistical significance. We discovered that functional modules in BrainRNNs are significantly neither merely localized nor randomly distributed (*p* < 0.001 for both Func1 in Fig. 4D and Func2 in Fig. S4B). A similar pattern was observed in the human brain, as demonstrated by the Yeo7 network (Fig. 4D) and the Yeo17 network (Fig. S4B). Notably, an analogous trade-off between spatial localization and distribution was also observed for network modules derived from structural connectivity in both BrainRNNs and the human brain (Fig. S4B). We further examined MWD across different wiring-cost constraint parameters λ (Fig. 4E). As expected, increasing wiring-cost constraints lead to a decrease in MWD for both functional and network modules, which means a more spatial localized module. These results indicated the complementary localized and distributed characteristic of functional modules are adjusted by structural constraints.

We then evaluated whether the functional modules are anchored by the predefined visual-motor-association segregation and network modules (Fig 4F). Overlap between partitions was quantified using adjusted mutual information (AMI), which enables comparison across different numbers of modules. Permutation tests by randomizing the labels across units were used to construct null distributions and assess statistical significance. In the sample model (the functional modules and the network module are displayed in Fig. 4B&C), functional modules showed significant overlap with network modules (Func1: AMI = 0.24, *p_perm_* < 0.0001; Func2: AMI = 0.20, *p_perm_* < 0.0001; *p* value calculated using permutation test denoted as *p_perm_*). A comparable relationship was observed in the human brain, where structural network modules exhibited significant overlap with functional modules (Yeo7: AMI = 0.21, *p_perm_* < 0.0001; Yeo17: AMI = 0.27, *p_perm_* < 0.0001). Moreover, both network and functional modular organization in BrainRNNs was significantly aligned with the predefined visual, motor, and association segregation, as indicated by significant AMI values between this pre-defined partition and the structural and functional modules (AMI = 0.36, *p_perm_* < 0.0001 for network modules, and AMI = 0.31, *p_perm_* < 0.0001, for functional modules). Together, these results indicate that functional modules in BrainRNNs are anchored by both areal visual and motor interface and formed network modularity.

A more critical question concerns how structural constraints give rise to this organization of functional modules. Previous studies have suggested that functional modularity in RNNs may emerge intrinsically for representing multiple tasks^21^. This implies that the functional modules can be decoupled from the structure without any structural priors. Here, we hypothesized that topographic structure anchors topological structure through the wiring-cost constraint, which further leads to the emergence and alignment of functional modules within the network. Wiring-cost constraints provide a mechanistic bridge linking structure and function. To test this hypothesis, we quantified both network modularity and functional modularity, quantified by the clustering coefficient, across all trained models with different wiring-cost constraint parameters (Fig. 4G). In the absence of wiring-cost constraints (λ = 0), functional modularity remained high. As wiring-cost constraint parameter λ increased, functional modularity remained stable and did not increase monotonically. By contrast, network modularity rose monotonically with increasing wiring-cost constraint parameter λ. We examined the correspondence between network modules and functional modules, quantified by their AMI (Fig. 4H). Notably, the overlap between network and functional modules increased monotonically with the increase of λ. Together, these results indicated that the structural constraints promote the emergence of topographically localized network modules which increase the modularity of topological structure and strengthens the coupling between network structure and functional modular organization.

To further explore the interrelationships among topography, topology, and functional modular organization, we examined the correlations among mean MWD of network modules, network modularity, and the AMI between network and functional modules (Func1). We found a strong negative correlation between MWD and network modularity (*r* = -0.92, *p* = 3.16 × 10^-49^, Fig. 4I), indicating that network modularity might arise from topographic constraints on the wiring-cost, which promote dense local connectivity within specific topographical regions. The coupling between functional and network modules was positively correlated with network modularity (*r* = 0.80, *p* = 3.22 × 10^-28^, Fig. 4J) and negatively correlated with the MWD of the network (*r* = -0.77, *p* = 9.77 × 10^-25^, Fig. 4K). Similar relationships exist when using Func2 (Fig. S4C&D). Together, these relationships indicated how topographic organization anchors network structure and, further construct the functional modular organization on the network topology. This chain might explain how prior studies can discover and segregate the functional modular organization based on both the topographic and topological structure without specific task-based activation.

### Discovering functional gradients on the structure

Brain function is distributed along functional gradients across the cortex^44^. The most prominent gradients are the sensorimotor-association and sensory-motor functional gradients (Fig. 5A), which can be recovered from intrinsic functional connectivity, and has strong agreement with the distribution of multiple structural factors, like myeline and cortical thickness^4,44^. To test whether gradients exhibit meaningful functional transition, we examined whether functional modules occupy distinct positions along the gradient using a Kruskal–Wallis (KW) test. As expected, Yeo7 functional modules were distributed non-uniformly along the gradient (Fig. 5B), with significantly distinct central tendencies across modules (*p* = 3.19 × 10^-70^, KW test). Moreover, topographic structure is reflected in the functional gradients. As previous study^44^, we calculated the minimal Euclidean distance between each cortical parcel and the 20 parcels with the highest gradient values, and correlated these distance with the principal functional gradient (*r* = -0.74, *p* = 2.62 × 10^-69^; Fig. 5C). These results indicate that spatial proximity along cortical coordinates is systematically related to functional differences captured by the gradient. Together, these discoveries suggest that functional gradients are shaped by both the topological and topographical structure.

**Fig. 5.**
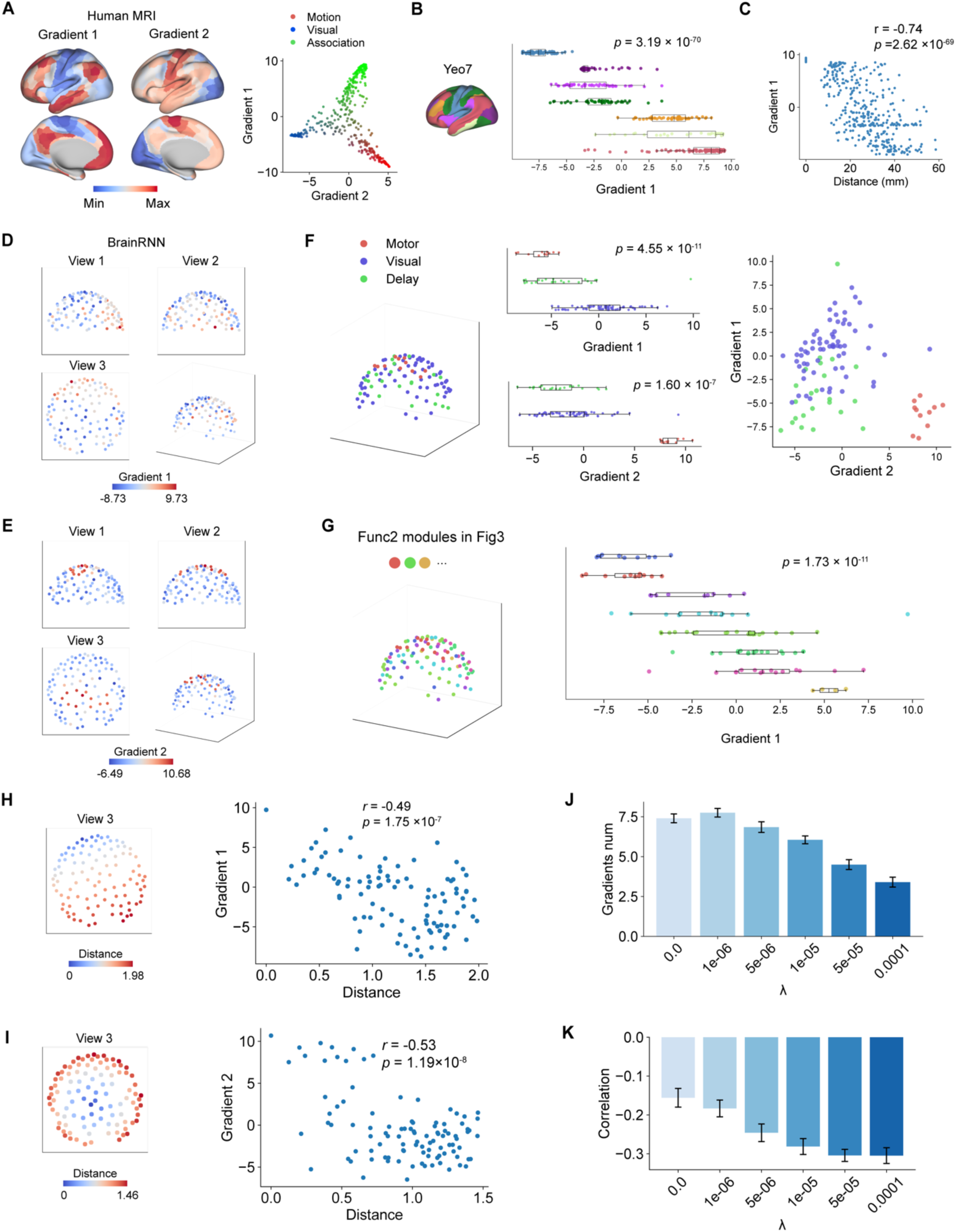
| Discovering functional gradients on the BrainRNN structure. **A**, The first and second functional gradients (Gradient 1&2) of the human brain, obtained from *BrainSpace* toolbox^45^. The first and second functional gradients represent sensorimotor-association and sensory-motor functional axes, respectively. **B,** Function modules in the human brain (Yeo7) are distributed along the first gradient, significantly locating at different gradient position. **C,** The first functional gradient is negatively correlated with the minimal Euclidean distance between each cortical parcel and the 20 parcels with the highest gradient values. **D-E,** The first and second gradients (Gradient 1&2) derived from the recurrent connectivity of the BrainRNN model. Units are displayed at their three-dimensional spatial coordinate and colored by gradient values. **F**, Func1 modules are significantly distributed along both the first and second gradients, with units preferentially activated for visual, motor, and delay periods occupying distinct gradient positions. **G**, Func2 modules occupy significantly distinct positions along the first functional gradients. **H-I**, Euclidean distance between each unit and the unit located at the highest position along the first (**H**) and the second (**I**) gradient. These distances are negatively correlated with the gradient values. **J**, The average number of significant functional gradients decreases with increasing wiring-cost constraint parameters λ. **K**, The average correlation between the first three gradients and distance to the unit with the highest gradient become more negative with increasing wiring-cost constraints λ. Error bars in **J**&**K** denote SEM.

To examine whether functional gradients can emerge from network structure in BrainRNNs similar to the human brain, we derived gradients by performing principal component analysis on the cosine similarity matrix of the recurrent connectivity. The first two gradients are visualized in three-dimensional views (Fig. 5D&E). To determine whether the components can be interpreted as functional gradients that encode functional transition, we projected the functional modules identified in the previous section (both Func1 and Func2, displayed in the Fig. 4B) onto these components (Fig. 5F&G). Based on the activation preference of Func1 for different periods in the task (Fig. 4B), we named the three modules as modules for visual, motor, and delay. We found the first and second components significantly encode the motor-delay-visual (*p* = 4.55 × 10^-11^, KW test) and delay-visual-motor (*p* = 1.60 × 10^-7^, KW test) gradient, mirroring the organization observed in the human brain (Fig. 5A). A finer-grained task-based functional modules (Func2) also exhibited significantly distinct positions along these functional gradients (*p* = 1.73 × 10^-11^, KW test, Fig. 5G for the first gradient; *p* = 4.04 × 10^-6^, KW test for the second gradient). These results demonstrate that functional gradients can be derived from the topological structure in the BrainRNN. To evaluate the topographic influence of the functional gradients, we calculated the Euclidean distance between each unit and the unit located at the highest position along each gradient axis (Fig. 5H&I for the first and second gradients, respectively). These distances exhibited significant negative correlations with gradient values (*r* = -0.49, *p* = 1.75 × 10^-7^ for the first gradient; *r* = -0.53, *p* = 1.19 × 10^-8^ for the second gradient), consistent with patterns observed in the human brain (Fig. 5C). This result indicates that functional differences captured by functional gradients were distributed along the underlying topographic structure.

Finally, we examined how structure constraints shape functional gradients. We quantified the number of significant functional gradients (*p* < 0.05 by KW test) across BrainRNNs trained with different wiring-cost constraints parameter λ (Fig. 5J). We found that the number of significant functional gradients decreased as λ increased. This reduction might partly account for the lower cognitive capacity observed under stronger wiring-cost constraints after training (Fig. 2D). Additionally, the average correlation between the first three gradient values and inter-unit distance increased with stronger wiring-cost constraints (Fig. 5K), indicating that functional gradients become more aligned with topographic structure under higher wiring-cost constraints. Together, these results suggest that structural constraints are accompanied by reduced functional diversity and stronger topographic embedding of functional gradients.

In summary, these results indicated that functional gradients capturing transitions across multiple functional domains could be discovered from topological structure and are distributed along the underlying topographic organization in BrainRNNs, similar to the discoveries in the human brain^44^. Moreover, structural constraints regulated the number of effective functional gradients supported by the network, which influence the computational capacity.

## DISCUSSION

In this work, we introduce brain-like recurrent neural network (BrainRNN), a recurrent neural network that integrates hemispheric topographic embedding, area-specific input and output segregation, and wiring-cost constraints into a comprehensive structural framework to mimic the human brain. We demonstrate that biologically motivated structure can shape connectivity, functional activation patterns, and cognitive capacity. Moreover, canonical principles linking structure and function in the human brain, including structure–function coupling, functional modules, and functional gradients, can also be applied to artificial neural networks. Moreover, by varying structural factors, we show that structural constraints enable functional organization to be inferred from structure. This may help explain why function is often interpreted through structural organization in neuroscience but less directly in artificial intelligence.

Linking structure to function is a central problem in neuroscience^7,8^. In the brain, multiple macroscale structural factors jointly provide the scaffold that supports and constrains the signal communication, which further shapes the cognitive function and behavior of the brain^36^. Structural features including white-matter pathways^36^, cortical geometry^26^, and spatial distance^27,39^, together with vasculature^46^, and hemodynamic dynamics^47^, have been shown to predict patterns of functional communication, which can further relate to inter-individual differences in cognitive function^8^. This linking allows clinical usage for understanding the causes and lead to discovery of biomarkers for a wide range of psychiatric and neurological disorders, including ADHD^48^, depression^9^, schizophrenia^49^, and Alzheimer’s^50^. By contrast, interpretability in artificial neural networks has largely focused on task representations and post hoc attribution of functional relevance to individual units, often through gradient-based methods^10,11^. Whether the structure– function linking approaches established in neuroscience can be extended to artificial systems remains unclear. Addressing this question requires explicitly constructing biologically meaningful structural organization, including topography and topology, in artificial networks, rather than relying on freely parameterized connectivity optimized solely for task performance. Here, we provide such a framework by integrating comprehensive macroscale structural constraints into artificial neural networks to simulate structurally constrained brain circuits. Our results replicate canonical findings in the human brain linking structure and function to demonstrate that these approaches can indeed be applied to the understanding of both biological brain and artificial intelligence.

The organization of the human brain is shaped by the interplay between functional integration across large-scale networks and functional segregation into specialized regions^1,2,5^. This organization gives rise to functional boundaries that are neither purely localized nor randomly distributed, characterized by functional parcellations^1,3^ and network modules^2^. BrainRNNs closely recapitulate these core organizational principles observed in the human brain. Moreover, by varying structural constraints, we further provide mechanistic evidence that macroscale structural constraints might contribute to the coupling between tasked-evoked functional segregation and structure-based segregation, enabling the discovery of function from the structure, as is commonly done in systems neuroscience. Beyond discrete modules, cortical function is also organized along gradients or hierarchies that capture transitions in functional specialization^4,44^. In particular, the sensorimotor–association hierarchy represents a dominant organizing axis that is not arbitrary but tightly constrained by diverse neurobiological properties, including anatomical structure and transcriptional profiles^4^. Our BrainRNNs support these observations by demonstrating that functional gradients can also be discovered from underlying network structure in artificial neural networks. Structural constraints allow functional gradients to be distributed along topographic axes and limit the number of effective functional gradients for cognitive capacity. These results provide the evidence of using functional gradients to understand the brain function and how structure constrains functional gradients. In addition, we found that the spatial localization of external input and output interfaces, specifically those targeting visual and motor regions, plays a critical role in shaping functional organization, a factor that has received relatively little attention in prior modeling studies. Together, this work bridges biological and artificial intelligence from a structural perspective and reveals a shared principle, that functional organization can emerge from and be systematically inferred from structure. Structural constraints enable functional organization to emerge in a manner that is intrinsically coupled to the topographic and topological structure.

Our results highlight that structural constraints lead to the expansion of association regions for higher-order cognitive functions. In BrainRNNs, tasks that require information maintenance and manipulation and flexible decision-making preferentially recruit units in association regions, whereas simpler visuomotor tasks can be supported by purely visual and motor units. Importantly, with areal visual and motor interface, increasing wiring-cost constraints selectively suppress activation and connectivity within association regions, leading to a pronounced decrease in higher-order cognitive performance. By contrast, conventional and spatial embedded RNNs^18^ do not reflect such disproportional expansion of association regions. This pattern suggests that the expansion of association regions may be driven by specific structural constraints, particularly the segregation of visual and motor interfaces into distinct cortical areas rather than globally distributed inputs and outputs. These findings parallel key observations from biological evolution^51,52^ and human development^4^. Across mammalian evolution, association cortex has undergone disproportionate expansion relative to primary sensory and motor areas^53,54^. Similarly, during human development, association cortices exhibit prolonged maturation trajectories^4,55^. Together, our results provide a mechanistic account that the expansion of association cortex might reflect a trade-off between increased cognitive capacity and structural constraints.

Artificial neural networks are currently widely used in neuroscientific studies. Trained through gradient-based optimization, these networks can develop task-specific representations and circuit-level solutions that support complex behaviors^56^. Prior studies have demonstrated alignment between representations in artificial neural networks and empirical neural data^14,17,57^. These comparative studies have provided valuable insights into the computational mechanisms underlying neural activity^16,21,57–59^ and biological processes such as development^59^. Despite these successes, artificial neural networks are typically parameterized models whose underlying architectures remain far from the structural organization of biological neural systems. Previous studies incorporate topographical features into the neural representation prediction and demonstrate that they contribute to the prediction accuracy in both visual and language cortex^19,60^, which highlights the importance of considering structural factors into the alignments. Comparatively few studies have examined how individual structural priors shape functional organization and computation^18,22,61^. One study demonstrates that RNNs trained with distance-dependent wiring constraints can exhibit similar topological feature as the brain network^18^. However, previous studies have not jointly incorporated multiple macroscale structural factors and multitask training into artificial neural networks, both of which are critical for simulating cortical functional organization. With this modeling framework, BrainRNNs help elucidate how multiple structural constraints give rise to functional organization observed in the brain. Moreover, they serve as in silico experimental platforms, enabling controlled tests of structural hypotheses that are difficult or impossible to address directly in biological systems.

This study has several potential limitations. First, the structural constraints implemented in BrainRNNs are necessarily simplified. The geometry of the cerebral cortex is far more complex than a hemispheric surface, forming a highly folded manifold that substantially increases cortical surface area within limited physical space. Such geometric complexity is known to influence patterns of functional organization^4^ but cannot be fully captured by the simplified spatial embedding used here. However, the present model is not intended to reproduce cortical geometry in full detail, but rather to test whether fundamental structure–function relationships can be recovered under minimal and controlled structural constraints. Second, functional organization in the brain emerges from the interaction between structural constraints and biological learning mechanisms, including genetically guided expression and activity-dependent plasticity. In contrast, BrainRNNs are trained using gradient-based optimization, which differs fundamentally from biological learning rules such as Hebbian plasticity. Although these biological learning mechanisms remain incompletely understood, their absence in the current framework limits our ability to fully assess how structural constraints interact with learning processes to shape functional organization. Third, the present study primarily uses human MRI data as a reference for evaluating structure–function relationships. Whether similar structural constraints exert comparable or distinct influences on functional organization across different species remains an open question. Comparative investigations across species may provide further insight into the generality and evolutionary relevance of the principles identified here.

Notwithstanding these limitations, this work represents a bridge between biological and artificial neural networks by demonstrating how brain-like structural constraints can shape functional organization and enable functional organization to be inferred from the structure in artificial neural networks, as commonly done in systems neuroscience. These findings highlight the importance of incorporating biologically grounded structural principles into computational models for future neuroscientific studies.

## METHODS

### BrainRNN

The “brain-like” recurrent neural networks (BrainRNNs) are constructed by embedding RNN units into a hemisphere coordinate system akin to the cortex, and assigning regional function for primary visual input and motor outputs (Fig. 1A). Specifically, the recurrent units are distributed quasi-uniformly across a hemispheric manifold. The location of units is defined using the method of Fibonacci lattices^37^. This construction prevents alignment artifacts and produces a near-uniform area coverage with low computational cost^37^. Specifically, assume 𝑛 = 128 units of a BrainRNN are distributed on a unit hemisphere surface with radius equal to 1. Each unit is indexed by 𝑖 = 1, …, 𝑛 with three-dimensional coordinates (𝑥_𝑖_, 𝑦_𝑖_, 𝑧_𝑖_). We defined the x axis as the left-right (L-R) axis, y axis as the posterior-anterior (P-A) axis, and z axis as the dorsal–ventral (D–V) axis. External visual input including angular location tuning and task cue was restricted to the visual units with 𝑛_𝑣𝑖𝑠_ = 32 at the posterior region, aligned with the primary visual cortex^62^. We define the visual units by calculating the nearest 32 points around the location (0,1,0). Motor readouts were taken from motor units with number 𝑛_𝑚𝑜𝑡_ = 16 at an anterior zone positioned slightly rostral to the middle line, analogous to the primary motor cortex. Thus, we define the units by calculating the nearest 16 points near the line connecting (-0.7, -0.25, 1) and (-0.7, 0.25, 1). We defined other units outside the visual and motor units as association units. This geometric arrangement enforces a posterior-to-anterior flow of information: visual signals enter through posterior units, undergo recurrent processing, and are subsequently decoded by anterior motor readout units. As a result, the architecture provides a spatially organized reference network that mirrors key aspects of brain anatomy and supports sensorimotor cognitive tasks.

Specifically, the updating of recurrent dynamics can be equated as:

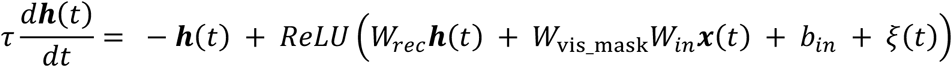

discretizing the updating function as

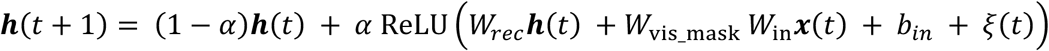

where ***h***(*t*) is the hidden state vector. *W*_rec_ 𝑎𝑛𝑑 *W*_in_denote the recurrent and input weight matrices. ReLU is used as the biologically inspired activation function that mimics neuronal thresholding mechanism. The hidden states of the motor units ***h***_motor_is linearly transformed to the predicted output vector as: **ŷ***_*t*_* = *W*_*out*_***h***_motor_ + 𝑏_*out*_, where *W*_*out*_ is the output weight matrix. All weights *W*_rec_, *W*_in_, *W*_*out*_ and biases 𝑏_𝑖𝑛_ 𝑎𝑛𝑑 𝑏_*out*_ are learnable parameters of the model. We set 𝛼 = 0.2. The *W*_𝑣𝑖𝑠_𝑚𝑎𝑠𝑘_is an identity positional matrix to project the input *W*_𝑖𝑛_𝒙(*t*) to the specific visual units. The vector 𝒙_*t*_ is the extrinsic visual input, which encodes the stimulus location cue (𝒙_loc_), fixation cue (𝑥_fix_), and task cue (𝒙_task_):

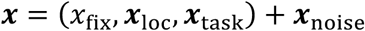

The fixation input is a binary signal indicating whether the subject was fixating (1) or not (0). The stimulus location input was encoded using a ring-like structure of 32 directionally tuned units that uniformly covered angles from 0 to 2𝜋. For a stimulus ***x***_𝑙𝑜𝑐_ with direction γ, the value of the *i* unit with preferred direction γ_i_was defined by:

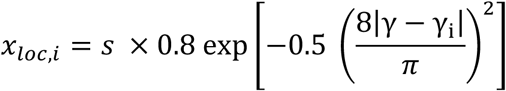

The task input is a one-hot vector with 22 units corresponding to the different task cue, respectively. Additive Gaussian noise ***x***_noise_ ∼𝓝(0, 0.01) was included in the input vector to simulate trial variability. Together, the fixation (1 unit), two stimulus modalities (32 units per modality), and task (22 units) components yielded a total of 87 input units.

The output **ŷ**_*t*_ was trained to predict the label 𝒚_*t*_, which includes the fixation output 𝑦_𝑓𝑖𝑥_, and the target saccadic direction 𝒚_𝑙𝑜𝑐_. The fixation output is greater than 0.85 before the response period and 0.05 during the response period. The target saccadic direction is encoded by the same 32-unit ring, identical to the input encoding. Each unit corresponds to a preferred γ_i_ ∈ [0,2π), and the target activity is defined as:

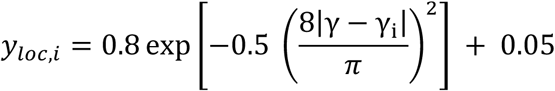

The response direction is read out from the population average response of the 32 unit direction^63^:

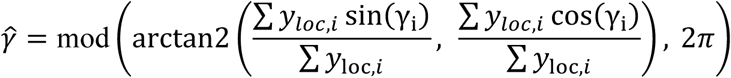

All initial input, recurrent, and output weights (*W*_𝑖𝑛_, *W*_𝑟𝑒𝑐_, *W*_*out*_) were drawn from 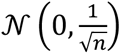. All biases are initialized to be zero.

### Training BrainRNNs with 22 cognitive tasks

We trained the model using 22 different cognitive tasks covering four different task families: visuomotor (VM), working memory (WM), decision-making (DM), and match-nonmatch (MNM) families. Across tasks, one or more stimuli can appear during the stimulus epoch, and depending on the rule, the model must: (i) produce an orienting response toward a direction on a ring, (ii) produce the opposite direction (anti), or (iii) repress any response (maintain fixation). Trials are divided into fix, stimulus (Stim1 or Stim2), delay (Delay1 or Delay2), and response epochs. The fixed cue is shown up in the fix, stimulus, and delay epochs, and might disappear in the response epochs. Depending on the task, one or two stimuli may appear during the stimulus epoch, with or without following a delay period. Across tasks, the model is either required to generate an orienting response toward an angular direction, to generate the opposite (anti) direction, or to repress any response by maintaining fixation. The following section will describe the task paradigm. The detailed time epochs and the stimulus mod and directions please see Table S1.

#### Visuomotor family

The visuomotor (VM) family implements the basic function for visual and motor of the BrainRNNs, including ‘Go’, anti-go (‘Anti’), reaction-time go (‘RT Go’), and reaction-time anti-go (‘RT Anti’), four tasks^21^. This family requires the plain information transportation from the visual units to the motor units. In the ‘Go’ task, a single stimulus appears early in the stimulus epoch and remains visible until the disappear of the fixation cue. Then in the response epoch, correct response is aligned with the stimulus direction (𝜃_target_ = 𝜃_stim_). In the ‘Anti’ task, stimulus and time set are same, except the required response is the opposite direction on the ring, 𝜃_target_ = (𝜃_stim_ + 𝜋) mod(2𝜋). The ‘RT Go’ task omits the fixation cue. The stimulus itself appears at the response epoch and serves as the imperative signal, so the model must respond immediately upon stimulus onset. The anti-version ‘RT Anti’ inverts the required direction in the same manner.

#### Working-memory family

In this family, besides the basic visual-motor information transportation, the networks are expected to maintain the information in their working-memory (WM), and also implements the cognitive control to filter the distractors^21,64^. The WM family includes delayed go (‘Dly Go’), delayed anti-go (‘Dly AntiGo’), remember-first (‘Rem1’), and remember-second (‘Rem2’). In the ‘Dly Go’ task, a delay is inserted between the stimulus and response epochs. The model must retain the direction through the delay and respond only after fixation offset. The anti-version ‘Dly AntiGo’ inverts the required direction in the same manner. The ‘Rem1’ and ‘Rem2’ tasks uses two sequential stimuli to test the maintenance of the directions and filter the other as the distractor. Specifically, two directions are presented sequentially within the same modality, with a delay afterward, and the required response after the final delay is to reproduce either the first (Rem1) or the second (Rem2) direction, respectively.

#### Decision-making family

The decision-making (DM) family presents two directions and asks the model to choose based on relative strength, with timing either simultaneous or separated by delays^65^, including decision-making (‘DM1’, ‘DM2’), contextual decision-making (‘Ctx DM1’, ‘Ctx DM2’), multisensory decision-making (‘MultiSen DM’), delayed decision-making (‘Dly DM1’, ‘Dly DM2’), contextual delayed decision-making (‘Ctx Dly DM1’, ‘Ctx Dly DM2’), and multisensory delayed decision-making (‘MultSen Dly DM’), ten tasks^21^. In the single modality DM1’ and ‘DM2’ task, both directions appear together during the stimulus epoch and persist until the response epochs. Target selection follows the stronger stimulus: 𝜃_target_ = 𝜃_stim1_if 𝑠_1_ > 𝑠_2_, and 𝜃_target_ = 𝜃_stim2_ otherwise, which 𝑠_1_and 𝑠_1_are the strengths for stim1 and stim2, respectively. In ‘ContextDM1’ and ‘ContextDM2’, both modalities carry stimuli, but the task input specifies which modality is behaviorally relevant. The same strength comparison applies for these two tasks, just restricted to the attended modality. In ‘MultiSen DM’, evidence is combined across modalities, and the target is chosen by the larger summed strength, 𝑠_1,mod1_ + 𝑠_1,mod2_versus 𝑠_2,mod1_ + 𝑠_2,mod2_. Delayed counterparts (‘Dly DM1’, ‘Dly DM2’, ‘Dly Ctx DM1’, ‘Dly Ctx DM2’, ‘Dly MultiSen DM’) present the two stimuli in separate time windows, with a delay epoch afterwards, requiring the model to maintain and compare evidence over time before responding at the end of the sequence.

#### Match-Nonmatch family

The match-nonmatch (MNM) family implements delayed match and non-match contingencies with either the directions or categories, including delayed-match-to-sample (‘DMS’), delayed-nonmatch-to-sample (‘DNMS’), delayed-nonmatch-to-category(‘DMC’), and delayed-nonmatch-to-category (‘DNMC’)^21,66^. In ‘DMS’ and ‘DNMS’, two directions are shown sequentially with an intervening delay. The second location cue determines whether to respond or to repress movement. In ‘DMS’, the model must produce a response only when two directions match; in ‘DNMS’, it must respond only when they do not match. When a response is required, the target direction is the direction of the second location cue in the DMS; otherwise, the correct behavior is to maintain fixation through the response epoch in the DNMS. Category-based variants follow the same structure but compare categories rather than exact directions. In ‘DMC’ and ‘DNMC’, directions are assigned to two categories by hemifield (e.g., [0, 𝜋) vs. [𝜋, 2𝜋)). Trials require a response if and only if the categories match for the DMC or do not match, for the DNMC. The response direction, when required, points to the direction of the second stimulus.

### Training models with wiring-cost constraints

The network was trained to minimize a loss function which minimizes the performance loss and wiring resources consumption. The loss of performance is 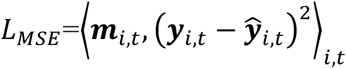, defined as the time-averaged mean squared error (MSE) between the predicted output **ŷ**_*t*_ and the target output 𝒚_*t*_, across output units *i* and time points *t*. The loss is weighted by a non-negative mask matrix 𝑚_𝑖,*t*_. During the stimulus and delay period, the ring output units has the mask weight equal to 1. In the response epoch, the first 100 ms was treated as a grace period with 𝑚_𝑖,*t*_ = 0, followed by an elevated weight for the response period 𝑚_𝑖,*t*_ = 5. For the fixation output unit, we used the 𝑚_𝑖,*t*_ = 2, to prioritize stable fixation control^21^. The loss of wiring-cost is defined as 𝐿_𝑐𝑜𝑛𝑛_ = 𝜆|*W*_𝑟𝑒𝑐_ ⊙ 𝐷|, where 𝐷 is the pair-wise Euclidian distance between units. We enumerate 𝜆 ∈ {0, 10^−6^, 5 × 10^−6^, 10^−5^, 5 × 10^−5^, 10^−4^} to evaluate the influence of wiring-cost constraints. Training was performed using batched gradient descent with a batch size of 128. The Adam optimizer was used with a learning rate of 0.001 and exponential decay rates of 0.9 and 0.999 for the first and second moment estimates, respectively. Model performance was evaluated based on the accuracy of responses during the response period. A response was considered correct if the decoded direction deviated from the target direction by less than 36 degrees. Each model was trained until all tasks reached at least 95% accuracy or 3,000,000 training epochs.

### MRI dataset and processing

#### HCP young adult (HCP) dataset

We analyzed multi-modal neuroimaging data from 297 unrelated participants (137 males, aged 22–36) from the HCP young adult (HCP) dataset (release S900), including T1-weighted structural, resting-state functional MRI (fMRI) and diffusion MRI^38^. Other details regarding the MRI parameters and our inclusive criteria have been described in our prior study^29^.

#### Structural and functional MRI data processing

Minimally preprocessed T1-weighted structural and functional MRI data were acquired from the HCP datasets^67^. The HCP minimal preprocessing pipelines used the tools FreeSurfer^68^, FSL^69^, and Connectome Workbench^70^. We then followed the post-processed protocols of eXtensible Connectivity Pipelines (XCP-D; https://xcp-d.readthedocs.io/en/latest/)^71^.

#### Functional connectivity construction with functional MRI

Functional connectivity (FC) refers to the temporal correlation of functional MRI signals. We computed FC by the following procedures. First, we extracted regional BOLD timeseries based on the prior Schaefer parcellation with 400 parcels^3^. Next, FC was calculated as the Pearson correlation coefficient between each pair of regional BOLD timeseries, resulting in a 400×400 symmetrical FC matrix for each participant. We then applied Fisher’s z-transformation to each FC value in the matrix for correlation analysis.

#### Calculating Euclidean distance between regions

With the T1w imaging, the centroid of each parcellation of the Schaefer400 template is calculated by averaging their coordinates. Then we calculated the Euclidean distance between two atlases by calculating the Euclidean distance between two centroids. This leaded to a 400x400 “Euclidean distance” matrix for each participant.

#### White matter structural network construction with diffusion MRI

We used minimally preprocessed diffusion MRI data from the HCP dataset^38^. The minimal preprocessing pipeline comprises b0 image intensity normalization across runs, EPI distortion correction, eddy current and motion correction, gradient nonlinearity correction, and registration to the native structural space (1.25 mm). The processed diffusion MRI data were further corrected for B1 field inhomogeneity using MRtrix3 (https://www.mrtrix.org/)^72^. We reconstructed whole-brain white matter tracts from preprocessed diffusion MRI data to construct the structural connectivity (SC). Reconstruction was conducted by the *mrtrix_multishell_msmt_ACT-hsvs* method in MRtrix3^72^, which implements a multi-shell and multi-tissue constrained spherical deconvolution (CSD) to estimate the fiber orientation distribution (FOD) of each voxel^73^. Then, we followed the anatomically constrained tractography (ACT) framework to improve the biological accuracy of fiber reconstruction ^74^. This tractography was performed by *tckgen*, which generates 40 million streamlines (length range from 30 to 250 mm, FOD power = 0.33) via a refined probabilistic streamlines tractography (iFOD2) based on the second-order integration over FOD. The streamlines were filtered from the tractogram based on the spherical deconvolution of the diffusion signal. We estimated the streamline weights using the command *tcksift2* ^75^. Next, the SC matrix was constructed by *tck2connectome* based on the Schaefer-400 atlas for each participant. The edge weight of SC indicates the number of streamlines connecting two regions. For each connection, we normalized the edge weight by dividing the average volume of the corresponding two regions^76^. Furthermore, the edge weight of the SC matrix was log-transformed, which is commonly used to shift the non-Gaussian SC distribution to the Gaussian distribution in previous studies^77^. Finally, this yielded a 400×400 symmetric SC matrix for each participant.

### Calculating structure-function coupling in the Human MRI and BrainRNNs

We evaluated the structure-function coupling using the Pearson correlation between structure profiles (Euclidean distance and the cosine similarity of structural connectivity) and functional connectivity for both human MRI data and BrainRNNs. The construction of Euclidean distance, structural connectivity, and functional connectivity in the human MRI is listed above in the MRI data processing procedure. As for BrainRNNs and other RNNs, the Euclidean distance matrix is the inter-distance matrix of the pre-defined coordinate of each unit. We defined the structural connectivity as the connection strength (absolute value of the connection weights) of the model. The functional connectivity is the temporal correlation of average unit responses across trials without the task cues (set the task cue input as 0), to simulate the resting-state functional connectivity. We extracted the upper triangle elements (79,800 unique edges) from either the 400×400 Euclidean distance or cosine similarity of structural connectivity matrices, creating a structural profile vector. Simultaneously, we extracted the upper triangle elements from the functional connectivity matrix, forming a functional profile vector. We calculated the Pearson correlation between the structural profile and functional profile across all connections at global level.

### Adjusted Mutual Information

We used adjusted mutual information (AMI) to quantify the overlap between two sets of discrete labels (modules). AMI measures the agreement between two partitions of the same set of units while correcting for the similarity expected by chance, allowing fair comparison across clustering solutions that differ in the number and size of modules^78^. Specifically, for two partition labels U and V, AMI is defined as

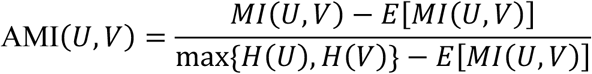

where 𝑀𝐼(𝑈, 𝑉) is the mutual information between the two labels, 𝐻(⋅) denotes the entropy, and 𝐸[𝑀𝐼(𝑈, 𝑉)] is the expected mutual information under a chance randomized assignment of labels. Thus, AMI = 1 indicates identical partitions while AMI = 0 corresponds to chance-level similarity. In our analyses, AMI was computed between the two modules over the same set of units. Permutation test was used to evaluate the significance of the AMI between two class. We preserved class counts while one of the labels were randomly permutated 𝐵 = 10000 times. AMI was recomputed for each permutation to form a null distribution, and a right-tailed 𝑝-value was estimated as (1 + #{AMI_perm_ ≥ AMI_obs_})/(𝐵 + 1).

### Mean within-module distance (MWD)

We utilized the mean within-module distance (MWD) to quantify the level of dispersion of modules across the hemisphere. Specifically, for a set of units 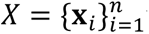 with module labels 𝑐 ∈ {1, …, 𝐶}, we first compute, for each module label 𝑐, the mean pairwise distance among its members:

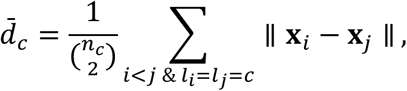

where 𝑛_𝑐_ is the number of samples in class 𝑐 and ∥·∥ denotes the Euclidean distance between two units. We then average the within-module distance across modules:

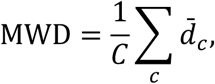

Lower MWD indicates tighter, more spatial compact module.

### Constructing localized and randomly distributed modules

To evaluate whether empirical modules exhibit both localized distributed and distributed across cortical regions, we constructed two types of theoretical module configurations: the pure localized modules and randomly distributed (no topographic constraint) modules. We aimed to evaluate whether the MWD of empirical modules are significant less than randomly distributed modules while greater than pure localized modules. To construct these two types of theoretical modules, we kept the number and size of empirical modules but redistributed the modules. An iterating searching algorithm is used to construct the localized modules. First, we initialized module centers by sampling the farthest points. One unit was randomly selected as the first center, and additional centers were iteratively chosen as the units that were farthest from the existing center set. This procedure continued until the overall number of module centers was selected. Second, each unit was assigned to a module using a nearest-center procedure. For every unit, we computed its distances to all module centers and sorted these distances in ascending order. Units were then greedily assigned to their closest available module until each module reached its target size. This step produced an initial partition that was spatially compact while exactly matching empirical module sizes. Third, we further optimized spatial compactness through an iterative swapping procedure. At each iteration, two different modules were randomly selected, and one unit from each module was sampled. The module labels of the two units were swapped if the swap led to a reduction in MWD. This procedure preserved module sizes while progressively decreasing spatial dispersion. The optimization terminated when no improving swap was found for 1000 attempts or when the number of iterations reached 10,000. The resulting modules yields spatially localized modules and exactly matching the prescribed module sizes. As a null baseline with identical modular size but no spatial structure, we generated randomly distributed modules by permutating module label. This preserves module sizes while removing any dependence on topography. To test the significance of whether the empirical modules are neither purely localized nor randomly distributed, we generated B = 1000 localized modules and random distributed modules for each empirical module. The *p* value is calculated as (1 + #{MWD_random_ ≤ MWD_obs_})/(B + 1) and (1 + #{MWD_local_ ≥ MWD_obs_})/(B + 1), respectively.

### Calculating network modules

To calculate the network modules for the structural connectivity in the BrainRNNs and human brain, community detection was performed using the Louvain algorithm as implemented in the *Brain Connectivity Toolbox*^43^. For the weighted network *W*_𝑖j_ ∈ 𝑾. The Louvain method maximizes the Newman–Girvan network modularity:

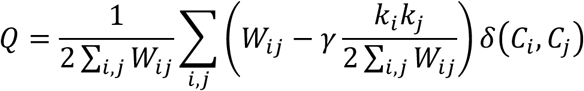

where 𝑘_𝑖_ = ∑_*j*_ *W*_𝑖*j*_. 𝛾 is the resolution parameter. Here we used default 𝛾 = 1. 𝛿(𝐶_𝑖_, 𝐶_*j*_) equals 1 when nodes *i* and *j* belong to the same community.

### Calculating functional modules

We identified functional modules by clustering units based on their task selectivity profiles^21^. For each task, the network was run across trials with a wide range of stimulus conditions. We computed the variance of each task across trials, measure for each unit by taking the variance of its activity across conditions at each time point excluding the fix period, averaged over time. The task variance was normalized by the maximal variance of all the tasks. Each unit’s normalized task variance values formed a feature vector representing its selectivity pattern. We applied k-means clustering to these vectors and evaluated modularity score using the silhouette score across solutions. To provide modules with different scale as the macroscale brain network^2,3^, a coarse-grained (2–5 clusters) and finer-grained modules (6–30 clusters) are calculated. The silhouette score represents the functional modularity used for other analysis.

### Calculating the functional gradients

Functional gradients in BrainRNNs were derived from recurrent network strengths. Following procedures commonly used to estimate functional gradients in the human brain^44^, we first computed the cosine similarity between the network strengths of all pairs of units. Principal component analysis (PCA) was then applied to the resulting similarity matrix. We assessed whether these principal components can be interpreted as meaningful functional gradients by testing whether different functional modules occupy distinct positions along each gradient, rather than being randomly distributed, using a Kruskal–Wallis (KW) test. As a comparison, macroscale functional gradients of the human brain were obtained from *BrainSpace* toolbox^45^.

## Data availability

The HCP datasets are available at https://db.humanconnectome.org/.

## Acknowledgements

We thank Mika Rubinov and Tatsuo Okubo for invaluable discussions, comments and feedback. This work was supported by NIH grant R01 EY036089. Data were provided [in part] by the Human Connectome Project, WU-Minn Consortium (Principal Investigators: David Van Essen and Kamil Ugurbil; 1U54MH091657) funded by the 16 NIH Institutes and Centers that support the NIH Blueprint for Neuroscience Research; and by the McDonnell Center for Systems Neuroscience at Washington University.

## Author contributions

P.C. designed the study, performed the modeling and analyses, and wrote the initial manuscript with review and editing from all other authors. C.C. supervised the study as the senior author. Z.C provided resources and instructions.

## SUPPLEMENTARY INFORMATION

**Fig. S1.**
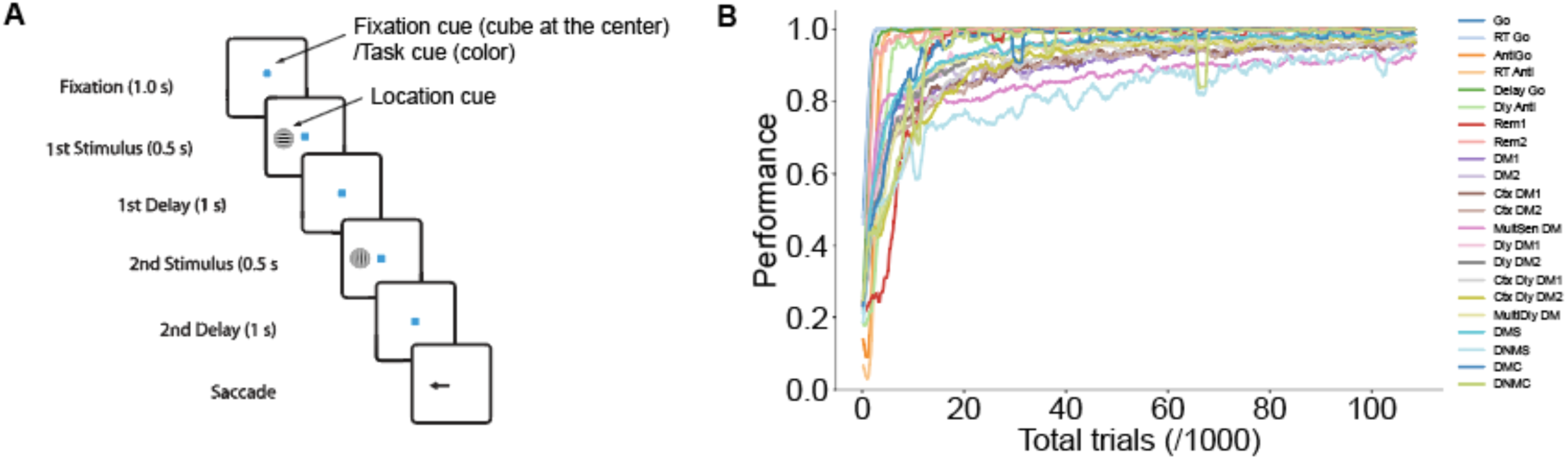
| Task scene and training. **a**, A sample task scene of the Remember-first task. **b**, Task performance during training of a sample model.

**Fig. S2.**
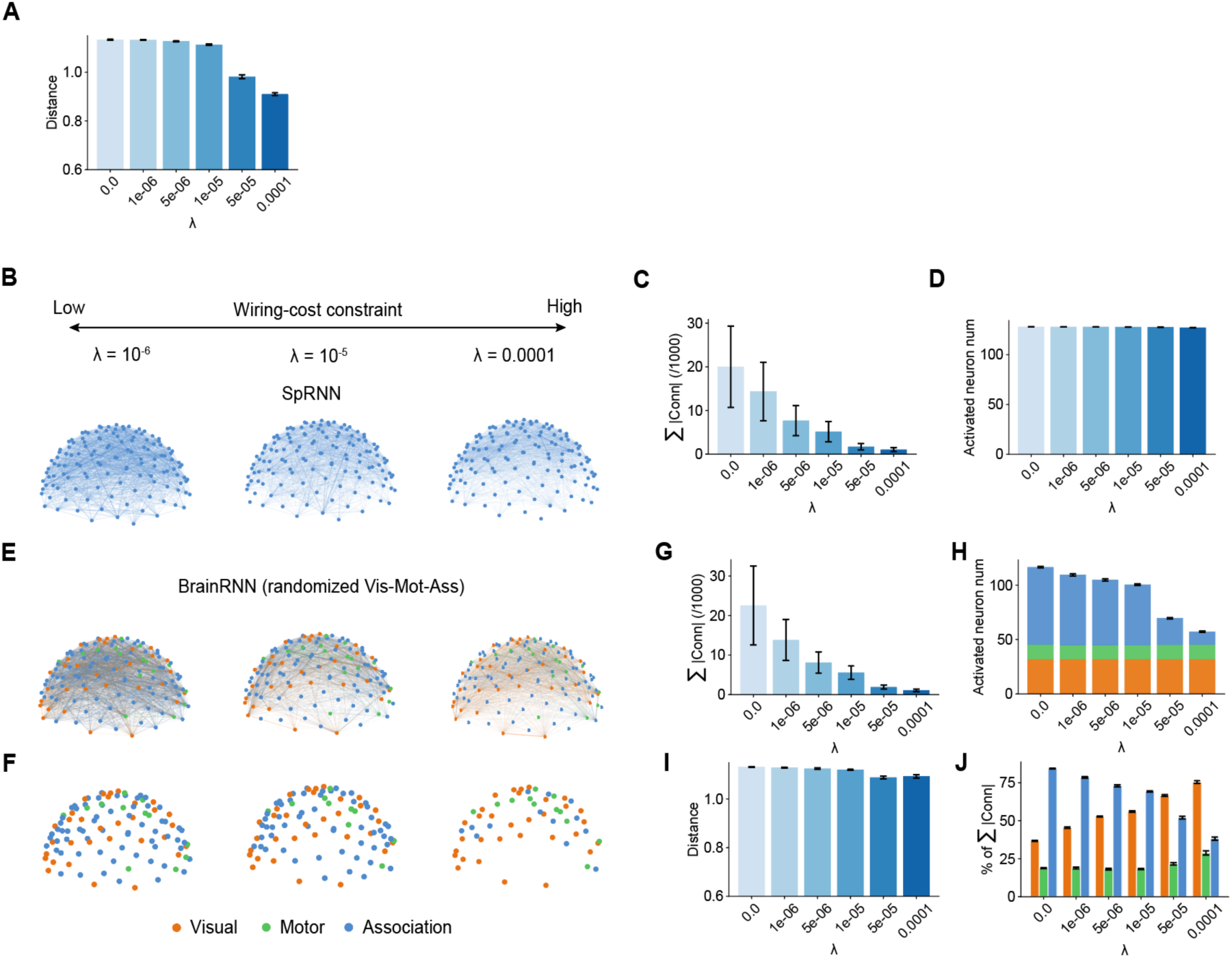
**A**, Average inter-units distance of activated units in BrainRNNs across wiring-cost constraint parameters. **B,** 3-Dimensional view of the spatial-embedded RNNs trained under different wiring-cost constraint parameters λ. The width of the line represents the connection strength (absolute value of connection weights). Top 10% absolute value of connection weights are displayed here. **C,** Mean total connection strength across Conventional RNNs (CRNNs, when λ = 0) and Spatial-embedded RNNs (SpRNNs, λ > 0) trained with different wiring-cost constraint parameters λ. **D**, Increasing the wiring-cost constraint parameters λ does not change the number of activated units in the conventional leaky RNNs. The visual-motor-association indices are randomized in the BrainRNNs for **E-J**. **E,** 3-Dimensional view of the BrainRNNs trained with randomized visual-motor-association unis trained under different λ. The width of the line represents the connection strength. Top 10% absolute value of connection weights are displayed here. **F,** 3-Dimensional view of the activated units. **G,** Mean total connection strength trained with different λ. **H**, Increasing the λ decreases the number of activated units, mainly in the association regions. **I,** Average inter-units distance of activated units in BrainRNNs with randomized visual-motor-association unis across different λ. **J,** Proportion of recurrent connection strengths associated with visual, motor, and association units.

**Fig. S3.**
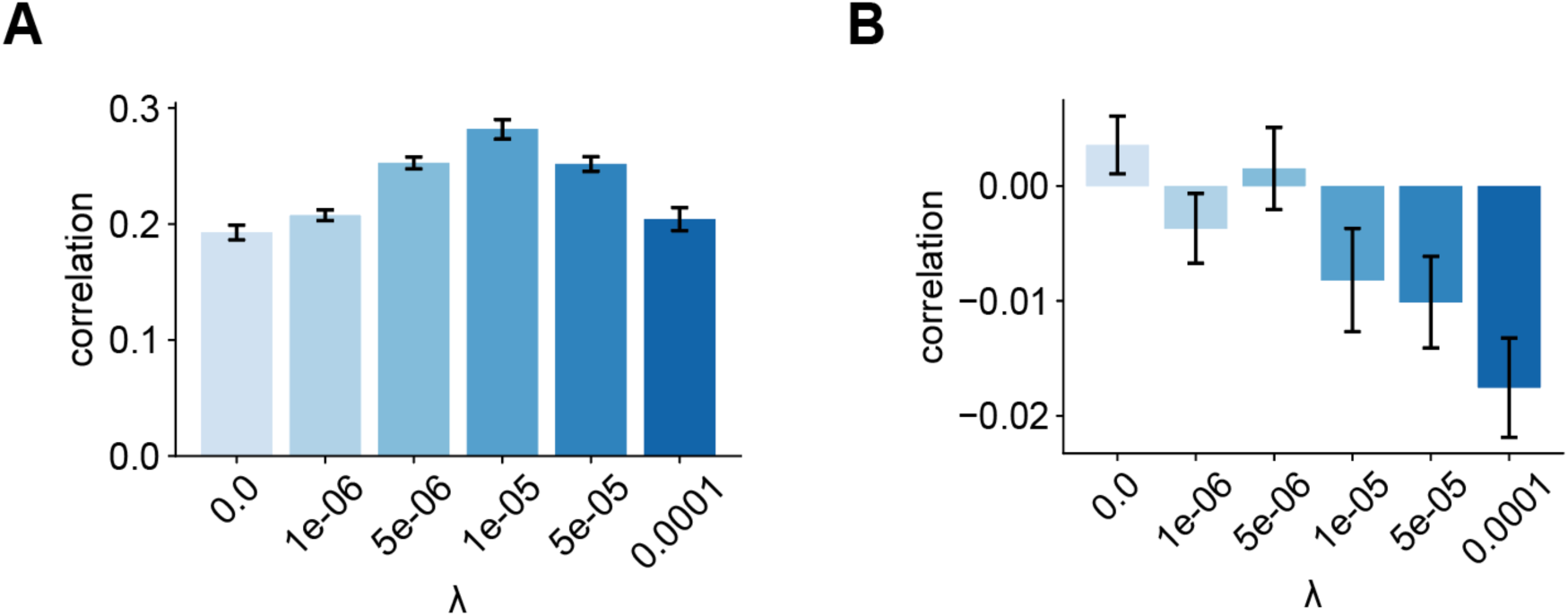
| Structure-function coupling in the conventional and spatial-embedded RNNs. **A-B,** Correlation between structural connectivity and functional connectivity (A) and between Euclidean distance and functional connectivity (B) in the conventional RNNs and spatial-embedded RNNs. Error bar denotes SEM.

**Fig. S4.**
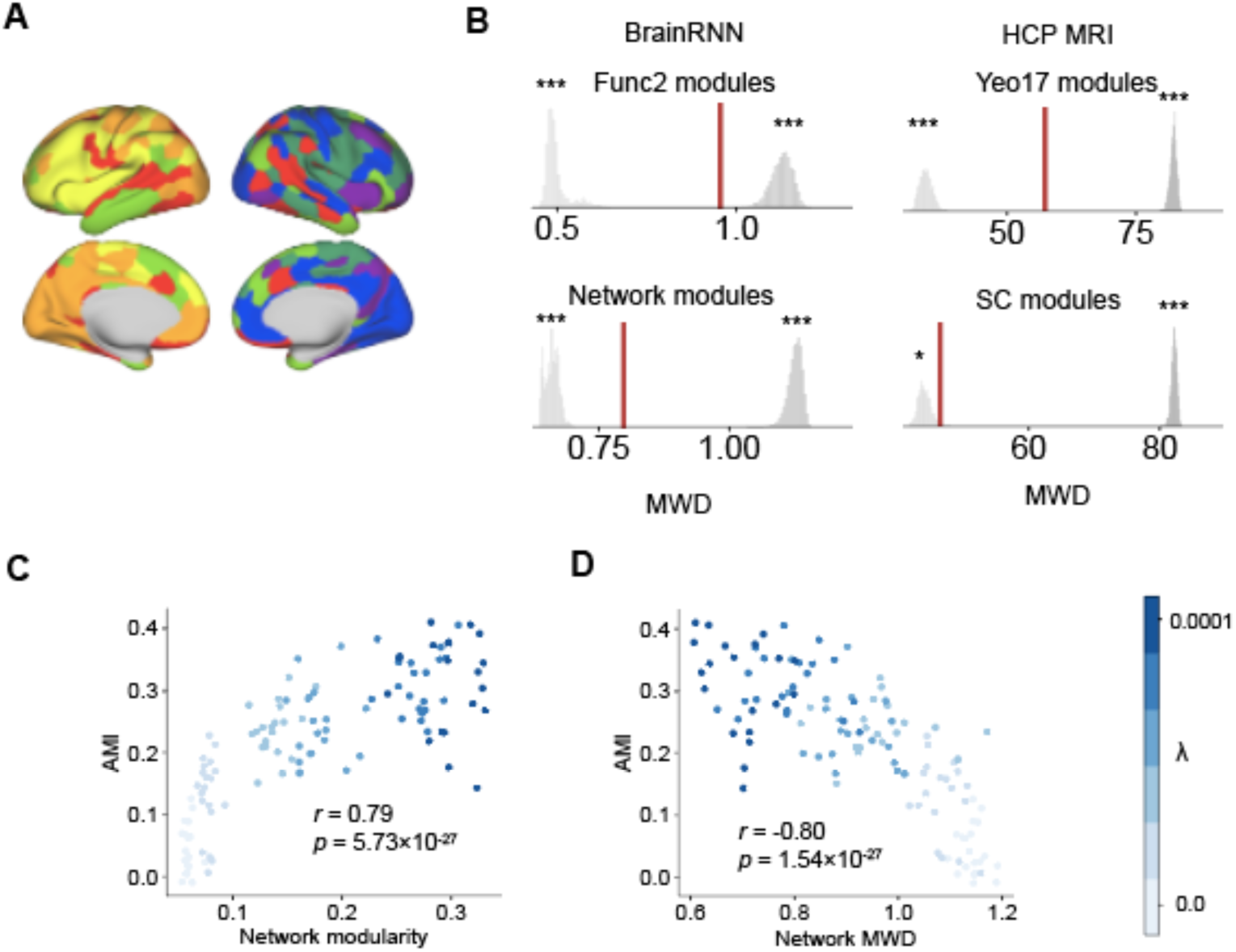
| **A,** Network modules in the human brain. **B,** Mean within-module distance (MWD) of BrainRNNs and human MRI data. **C**-**D**, Correlations between modularity of network modules (**C**), MWDs of network modules (**D**) with AMI between functional (Func2) across BrainRNNs trained with different wiring-cost constraint parameters λ. * denotes *p* < 0.05 and *** denotes *p* < 0.001.

